# Microglial TLR4 Mediates Post-UTI Chronic Pelvic Pain

**DOI:** 10.64898/2026.08.04.742321

**Authors:** Habib Jmii, Shivesh Ghura, Anthony J. Schaeffer, David J. Klumpp

## Abstract

Urinary tract infections (UTIs) are a major risk factor for interstitial cystitis/bladder pain syndrome (IC/BPS), yet the mechanisms driving chronic pelvic pain and associated symptoms remain poorly understood. Here, we investigated the role of microglia and Toll-like receptor 4 (TLR4) in a mouse model of post-UTI chronic pelvic pain (PUPP). Infection with *E. coli* induced persistent pelvic allodynia that was significantly attenuated by microglial depletion (PLX5622) or inhibition (minocycline), indicating a key role for microglia in pain maintenance. In contrast, microglial depletion did not improve urinary dysfunction or anxiety– and depression-like behaviors. Prefrontal cortex microglia of PUPP mice exhibited reduced microglial branching complexity and a less ramified phenotype, indicative of an activated microglial state. Transcriptomic profiling of brain CD11b+ cells revealed a reactive microglial signature enriched for chemokines, NFΚB-related genes, and immediate early response genes, alongside pathways involved in immune regulation and leukocyte recruitment. Both general and microglia-specific TLR4 deletion reduced pelvic allodynia and reduced microglial morphological features of activation. Consistent with this, pharmacological TLR4 inhibition in vitro suppressed LPS-induced NFΚB activation, cytokine secretion, and CD68 expression. Together, these findings identify microglial TLR4 as a critical mediator of post-UTI chronic pelvic pain.

## Introduction

Interstitial cystitis/bladder pain syndrome (IC/BPS) is a chronic disorder characterized by pelvic pain, urinary urgency, and increased urinary frequency, affecting nearly 12 million people in the United States **[1–3]**. IC/BPS is also commonly associated with anxiety and depression, significantly reducing quality of life **[4,5]**. A history of urinary tract infection (UTI) is commonly reported among patients with IC/BPS, raising infection as one proposed etiology for IC **[6–9]**. Acute UTI disrupts urothelial integrity and causes bladder inflammation and sensory nerve responses **[9–11]**. In contrast IC/BPS symptoms are chronic, suggesting involvement of central nervous system (CNS) mechanisms. Central sensitization is increasingly recognized as a major driver of chronic pelvic pain, where persistent nociceptive signaling induces long-lasting amplification of CNS pain pathways **[12–15].** Consistent with this, neuroimaging studies in IC/BPS patients have demonstrated altered brain activity associated with pain severity **[16–18]**.

Microglia, the resident immune cells of the CNS, are key regulators of neuronal plasticity and neuroinflammation and contribute to chronic pain through the release of pro-nociceptive mediators and modulation of synaptic activity **[19–22]**. Among the pathways involved in microglial activation, Toll-like receptor 4 (TLR4) has emerged as an important mediator of various types of pain in both peripheral organs and central nervous system. TLR4 signaling promotes neuroimmune activation and pain sensitization, and studies have shown that TLR4 inhibition or deletion reduces pain hypersensitivity in multiple pain models **[23–25]**. In IC/BPS, clinical and experimental studies, including findings from the NIDDK MAPP Research Network, have associated TLR4-related inflammatory signaling with symptom severity and chronic pain **[26,27]**. However, these studies have primarily focused on peripheral mechanisms, while the contribution of CNS TLR4 to IC/BPS remains unknown.

In experimental UTI in mice, we have previously shown that acute infection with *E. coli* can elicit a spectrum of pain behavior responses, ranging from acute pain to post-UTI chronic pelvic pain or even analgesia, depending upon the clinical origin of strains and the status of O-antigen **[28]**. Indeed, strains lacking O-antigen cause post-UTI chronic pelvic pain (PUPP), that persists long after bacterial clearance and that is associated with central sensitization **[28,29]**. Because IC/BPS patients often have a history of prior UTI but where a diagnosis that excludes current UTI, PUPP is a model for the postulated infectious etiology of IC **[30]**. Here, we demonstrate that chronic pelvic pain in the PUPP model is associated with microglial activation and depends on TLR4 signaling. Pain is modulated by targeting microglial activation, suggesting new therapeutic approaches for IC.

## Materials and methods

### Animals

Female C57BL/6J mice and *Tlr4* KO mice (B6(Cg)-*Tlr4tm1.2Karp/J*) were purchased from The Jackson Laboratory (JAX) and housed at Northwestern’s Center for Comparative Medicine on a 12h:12h light: dark cycle with unrestricted food or water. Mice were maintained and used under Northwestern IACUC-approved protocols and followed NIH guidelines to minimize pain or stress. Microglia-specific *Tlr4* conditional knockout (cKO) mice were generated by crossing *Tmem119-Cre* mice (JAX, C57BL/6-*Tmem119em1*(*Cre*/*Ert2*) *Gfng/J*) with *Tlr4* floxed mice (JAX, B6.129-*Tlr4tm1.1Jke/J*). The resulting offspring were backcrossed with *Tlr4* floxed mice to generate mice that carry the *Tmem119-Cre* transgene and homozygous for the floxed *Tlr4* allele (*Tmem119-Cre*; *Tlr4* flox/flox). The Cre/loxP recombination was induced by tamoxifen (75mg/Kg) injected intraperitonially for 5 consecutive days, and those mice were used as microglia-specific *Tlr4* cKO mice, while Cre-negative littermates served as controls.

### Induction of PUPP model

*E. coli* NU23 (NU23) is a clinical isolate of *E. coli* obtained from a patient with acute UTI **[31]**. NU23 was cultured overnight in Luria broth at 37°C with shaking followed by culture under static conditions for two days to promote the expression of type 1 pili (1:1000 dilutions). 10ul of NU23 suspension (10^10^ CFU/mL in PBS) were used for non-reflux transurethral instillation of eight– to twelve-weak-old female mice, performed under anesthesia using isoflurane.

### Pelvic allodynia

Allodynia was quantified in response to Von Frey filament stimulation to the pelvic region **[32]**. Mice were placed individually in test chambers and allowed to adapt to the test chamber environment for 10 minutes. Then, five Von Frey filaments of increasing force (0.04 to 4g) were applied 10 times each to the pelvic region, at distinct sites to avoid wind up, and responses (jumping, flinching, licking) were recorded. The response percentage, normalized to baseline, was calculated using the following formula: response change %= (sum responses-sum baseline responses/sum baseline responses) * 100.

### PLX5622 and minocycline treatments

PLX5622 and minocycline were used for microglia ablation or inhibition, respectively. PLX5622 (Med Chem Express, HY-114153) was dissolved in 10% DMSO and 90% corn oil. PLX5622 solution was then gavaged to mice (90mg/Kg) for 5 consecutive days to ensure maximum elimination of microglia. Minocycline, a tetracycline known for its capacity for inhibiting microglia activation **[33]**, was gavaged to mice (60mg/Kg) for 5 consecutive days. Pelvic pain, voiding activity, and anxiety and depression behaviors were assessed one day after PLX5622 or minocycline treatments. Then, drug administration was ceased to allow microglia repopulation and regain of activity. Ten days after drug administration cessation, pelvic pain was reassessed to investigate the role of microglia in post-UTI chronic pelvic pain.

### Assessment of voiding activity

The voiding activity was assessed by the void spot assay **[34]**. Mice were placed on an absorbent 3 MM filter paper for 90 minutes without water or food. Urine spots were then visualized and captured using Invitrogen iBright Imaging System (FL1000) by exposing the filter paper to UV light. The number of spots were counted and the volume per spot was determined by measuring the approximate area of the void spot in ImageJ and interpolating it against a calibration curve of water spotted onto filter paper.

### Anxiety and depression-like behavior assessment tests

#### The dark-light box assay

The dark–light box test is a behavioral assay used in rodents to measure anxiety-like behavior. Mice were placed in the lit side of the box and allowed to move freely between the two compartments (dark and light) for 10 minutes. Time in light chamber, latency to enter dark chamber, and the number of transitions were recorded to assess anxiety levels in studied mice. Animals naturally prefer dark enclosed spaces, but they also explore new environments. More anxiety-like behavior usually means the mouse avoids the bright chamber.

#### Novelty-suppressed feeding assay

Novelty-suppressed feeding test was performed to quantify depressive-like behavior. Mice were deprived of food for 24 h and water for 1h prior to testing. For the test, a food pellet was placed in the center of a large square chamber, and a mouse was then placed into a corner of the chamber, and time to approach and consume the food pellet were recorded using a stopwatch.

### Immunohistochemistry (IHC) and microglia skeleton analyses

Frozen brain sections were prepared for IHC as follows: mice were transcardially perfused for 2 min with phosphate buffered saline (PBS) followed by 4% paraformaldehyde for 10 min under anesthesia. The harvested brains were further fixed in 4% paraformaldehyde overnight and sequentially dehydrated in 15% and 30% sucrose solutions. Tissues were then frozen in dry ice and embedded in Tissue Plus optimum cutting temperature medium (Fisher HealthCare, Houston, TX) before storing at –80°C. Afterwards, cryostat sections of 20μm were generated and mounted on charged slides. Prior to immunostaining, sections were washed twice in PBS and incubated for 1 hour at room temperature with 5% bovine serum albumin blocking solution prepared in PBS. Sections were then incubated overnight at 4°C with the primary antibodies: goat monoclonal anti-Iba1 (Abcam, 289874, 1:400) and/or rabbit polyclonal anti-TLR4 (Novus Biologicals NBP2-24821, 1:250) diluted in normal antibody diluent (Skytech, ADT500). Brain sections were then washed 4 times with PBS and incubated for 90 min at RT with the secondary antibodies diluted as follows: Alexa Fluor 488-donkey anti-rabbit (Invitrogen, A-21206, 1:400) and Alexa Fluor 594-donkey anti-goat (Invitrogen, A-32758, 1:400). Secondary antibodies were then washed off using PBS and coverslips were mounted onto slides with antifade mounting medium with DAPI (ThermoFisher, P36971). Afterwards, slides were imaged using Nikon Eclipse Ti2 microscope with NIS-Elements imaging software. The morphological characterization of microglia was performed on images of the prefrontal cortex region from 4 different mice with a total of 15 fields analyzed as detailed in Young & Morrison **[35]**. ImageJ software and plugins (brightness/contrast, unsharp mask, despeckle, binary, close, remove outliers, and skeletonize) were utilized to convert images to binary and skeletonized images. The Analyze Skeleton Plugin was then applied to the skeletonized image which tags and measures microglia ramification including branches, endpoints, average process length, and longest process length. The data was then transferred to an Excel spreadsheet and organized by end-point voxels followed by maximum branch length (highest to lowest). Using the straight-line measure tool in Image J, outliers and noise were measured and excluded from the data set prior to summation of the data. In addition, cell somas were manually counted and summed data were divided by number of cells/image to quantify average morphology per cell.

### Microglia isolation and bulk RNA-sequencing

Microglia were isolated from the prefrontal cortex (PFC) of mice 3 weeks after bladder instillation with either PBS or NU23. The PFCs were minced using a scalpel then enzymatically dissociated using Miltenyi neural tissue dissociation kit as instructed by manufacturer (Miltenyi Biotec, 130-094-802). Afterwards, mononuclear cells were separated from the cell suspensions using isotonic Percoll gradient (30%-70% in PBS) and separated mononuclear cells were then incubated with CD11b magnetic beads (Miltenyi Biotec, 130-097-142) to isolate CD11b positive cells encompassing microglia. After magnetic separation of CD11b positive cells, cells were washed and resuspended in 500 ul of Trizol and RNA was extracted using the Direct-zol ^TM^ RNA MiniPrep Plus kit (Zymo Research, R2070) as instructed by the manufacturer. RNA-seq was conducted in the Northwestern University NUSeq Core Facility. RNA quantity was determined via Qubit fluorometer. Total RNA examples were also checked for fragment sizing using a Bioanalyzer 2100 (Agilent). The Illumina Stranded Total RNA Library Preparation Kit was used to prepare sequencing libraries from 10 ng of total RNA samples, according to manufacturer’s instructions. This procedure includes rRNA depletion with RiboZero Plus, cDNA synthesis, 3’ end adenylation, adapter ligation, library PCR amplification and validation. The Illumina NovaSeq X Plus sequencer was used to sequence the libraries with the production of single-end, 50 bp reads at the depth of 20-25 M reads per sample.

### Cells

BV-2 murine microglia cell line was purchased from AcceGen (ABC-TC212S). Cells were maintained in high-glucose DMEM supplemented with 10% fetal bovine serum (FBS), 1% L-glutamine, and 1% penicillin-streptomycin. To develop a microglia NFΚB reporter cell line, BV2 cells were transfected with pNiFty2-N-Fluc-Zeo plasmid (InvivoGen, pnf2-fluc) using lipofectamine 3000 as recommended by the manufacturer (ThermoFisher Scientific, L3000008).

Two days after transfection, successfully transfected cells were selected with zeocin (ThermoFisher Scientific, J67140.XF) applied at 1mg/mL for 10 days.

### TLR4 inhibition and cytokines measurement

To study the role of TLR4 in microglia activation, 10^5^ BV2 cells/well were plated in 12-well plates and incubated overnight at 37°C 5% CO2. Cells were then treated with TLR4 inhibitor TAK-242 (Sigma-Aldrich, 614316) at 5μM (final concentration) followed by a stimulation with LPS (Sigma-Aldrich, L4524) applied at 100ng/mL for 6h. Cell cultures supernatants were then collected and used for cytokine quantification by ELISA. We used R&D Systems mouse DuoSet ELISA kits to quantify TNFα (DY410) and IL-6 (DY40605) as instructed by the manufacturer.

### NFΚB reporter assay

Reporter activity was quantified using luciferase reporter assay. Briefly, TLR4 inhibitor TAK-242 (Sigma-Aldrich, 614316) or medium were added to NFΚB stably transfected BV2 cells (5μM) and incubated for 1h. Cells were then treated with a 100ng of LPS (Sigma-Aldrich, L4524) for 6h to induce NFΚB expression. After incubation with LPS, media were aspirated and cells were lysed, using 50 μL of passive lysis buffer (Promega, E1941), for 10 min at RT before adding 50ul of firefly D-luciferin solution and immediately measuring bioluminescence using Cytation3 plate reader (BioTek).

### TLR4 inhibition and IF for CD68

The increased expression of CD68 is a marker a microglia activation. Hence, we assessed the effect of TLR4 inhibition on CD68 expression in BV2 microglia. Cells were plated on poly-l-lysine coverslips placed in 12-well plates at a density of 10^5^ cells/well and incubated overnight. The next day, cells were treated for 1h with TLR4 inhibitor TAK-242 at 5μM before stimulating them with LPS at 100ng/mL for 6h. Cell culture media were then aspirated, cells were washed with PBS, and fixed using 4% PFA for 15 min at RT. Then, cells were washed with PBS, permeabilized for 10 min with 0.05 % Triton X-100 (Sigma Aldrich, X100), and non-specific binding sites were blocked using 5% BSA for 1h at RT before adding rabbit anti-CD68 antibody (Cell Signaling,

97778, 1:200), diluted in normal antibody diluent (Skytech, ADT500), and incubation overnight at 4°C. Afterwards, cells were washed 5 times with PBS and incubated for with Alexa Fluor 488-donkey anti-rabbit (Invitrogen, A-21206, 1:400) secondary antibody for 1.5h at RT. Secondary antibody solution was then washed off using PBS and coverslips were mounted onto slides using antifade mounting medium with DAPI (ThermoFisher, P36971). The slides were subsequently imaged using Nikon Eclipse Ti2 microscope with NIS-Elements imaging software.

The expression of CD68 was determined by quantifying fluorescence intensity using FIJI software as described by Shihan et al **[36]**.

## Results

### Microglia differentially contribute to IC-like symptoms

To establish a model of post-UTI chronic pelvic pain and investigate the mechanisms underlying persistent pain following urinary tract infection, we utilized the O-antigen-deficient *E. coli* strain NU23. O-antigen-deficient *E. coli* strains have been previously shown to induce chronic pelvic pain despite resolution of the acute infection **[28]**. While modest increases in pelvic sensitivity were observed one week post-instillation, NU23-instilled mice developed progressive pelvic allodynia over 3 weeks. A marked increase was evident by 3 weeks, resulting in significantly greater responses compared with PBS-instilled controls (p < 0.001). These findings indicate that infection with NU23 induces persistent pelvic pain that becomes fully established during the chronic phase after infection **(Fig. 1A)**. To investigate the contribution of microglia to pelvic pain, PUPP mice were treated with either the CSF1R inhibitor PLX5622 to deplete microglia or with vehicle. We found that PLX5622 treatment depletes approximately 80% of microglia **(Fig. S1**). Pelvic sensitivity was assessed one day after PLX5622 treatment and again 10 days after treatment cessation. PLX5622 treatment significantly reduced pelvic allodynia compared with vehicle-treated mice **(Fig. 1B)**. However, pelvic allodynia re-emerged after discontinuation of PLX5622 **(Fig. 1B)**. To further evaluate the contribution of microglia to NU23-induced pelvic pain, mice with post-UTI chronic pelvic pain were treated with the microglial inhibitor minocycline **[30]**. Minocycline administration produced a marked reduction in pelvic allodynia compared with pretreatment levels (*p < 0.05). However, this effect was not sustained, as pelvic allodynia increased following treatment cessation and approached levels observed in vehicle-treated mice **(Fig. 1C)**. Together, these findings suggest that pharmacological ablation or inhibition of microglial activity transiently alleviates post-UTI chronic pelvic pain.

**Figure 1.**
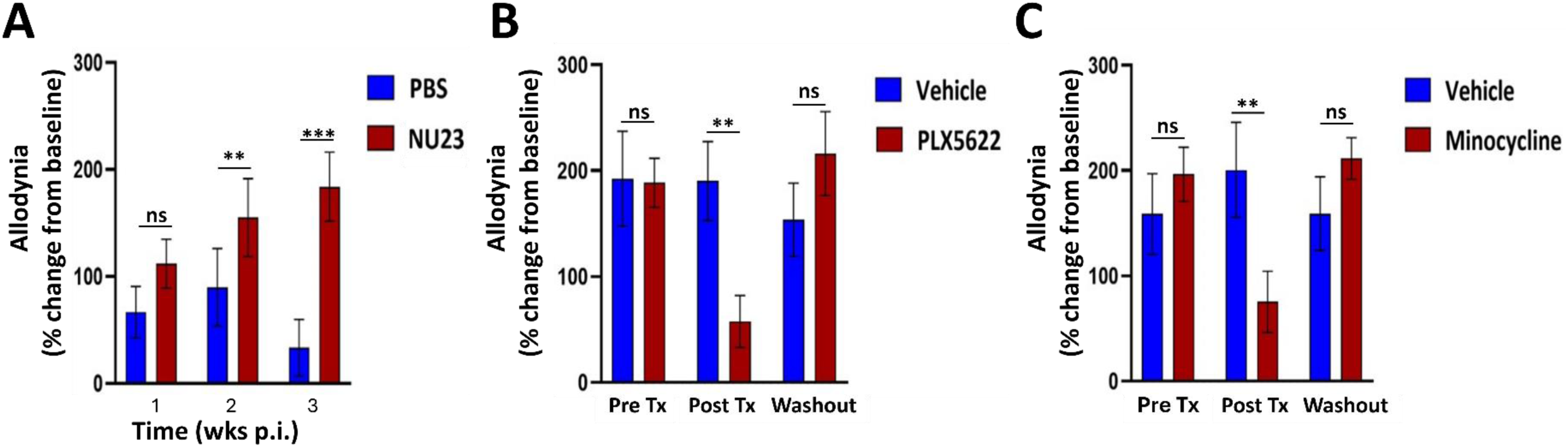
Microglial ablation or inhibition reverses established post-UTI pelvic pain. **(A)** Pelvic allodynia was assessed weekly in mice following transurethral instillation with PBS or *E. coli* NU23. Response frequency to Von Frey filaments stimulation was measured at week 1, 2, and 3 post-instillation and normalized to baseline. **(B)** Pelvic allodynia in NU23-instilled mice measured at 3 weeks post-instillation, following treatment with vehicle or microglia suppressor PLX5622 for 5 days, and 10 days after PLX5622 washout. **(C)** Pelvic allodynia in NU23-instilled mice measured at 3 weeks post-instillation, following treatment with vehicle or microglia inhibitor (minocycline) for 5 days, and 10 days after cessation of minocycline treatment. Data are presented as mean ± SEM. Statistical significance was determined using Welch’s t-test. *P < 0.05, **P < 0.01, ***P < 0.001.

To determine whether microglia contribute to urinary dysfunction associated with PUPP, voiding behavior was assessed following infection with NU23 and subsequent microglial ablation. Consistent with the development of PUPP, NU23 infection resulted in significant voiding urinary dysfunction characterized by increased urinary frequency. Similarly, NU23 infection causes an elevation of anxiety– and depression-like behaviors as demonstrated by novelty-suppressed feeding and light-dark box assays **(Fig. S2 A and Fig. S3 A, C)**. However, no significant differences were observed between microglia-depleted, using PLX5622, and control mice in voiding frequency and urine output, indicating that microglial depletion did not improve urinary dysfunction (**Fig. S2 B, D**). Similarly, assessment of anxiety– and depression-like behaviors revealed no significant effects of microglial ablation on performance in the novelty-suppressed feeding and the light-dark box tests **(Fig. S3 B, D)**. Together, these findings suggest that, unlike pelvic pain, urinary and affective symptoms associated with IC/BPS are independent of microglial signaling.

### PUPP is associated with microglial morphological remodeling

To determine whether chronic pelvic pain following NU23 infection is associated with alterations in microglial activation status, we performed morphological analyses of microglia in the prefrontal cortex, a critical region for chronic pain perception **[37]**. Briefly, Iba1-immunolabeled microglia were imaged and processed using Fiji/ImageJ, where cells were skeletonized to quantify branching parameters including total branch length, number of endpoints, and branch junctions. Morphological analysis revealed marked alterations in microglial architecture in NU23-instilled mice in mice with PUPP compared to with PBS the control ones. Specifically, microglia from NU23-instilled mice exhibited significant reductions in total branch length, number of endpoints, number of branch junctions, and overall ramification complexity **(Fig. 2)**. Such changes are indicative of a shift from a ramified homeostatic phenotype toward a more activated microglial state **[38]**. These morphological findings provide evidence of microglial activation in the brains of mice with PUPP.

**Figure 2.**
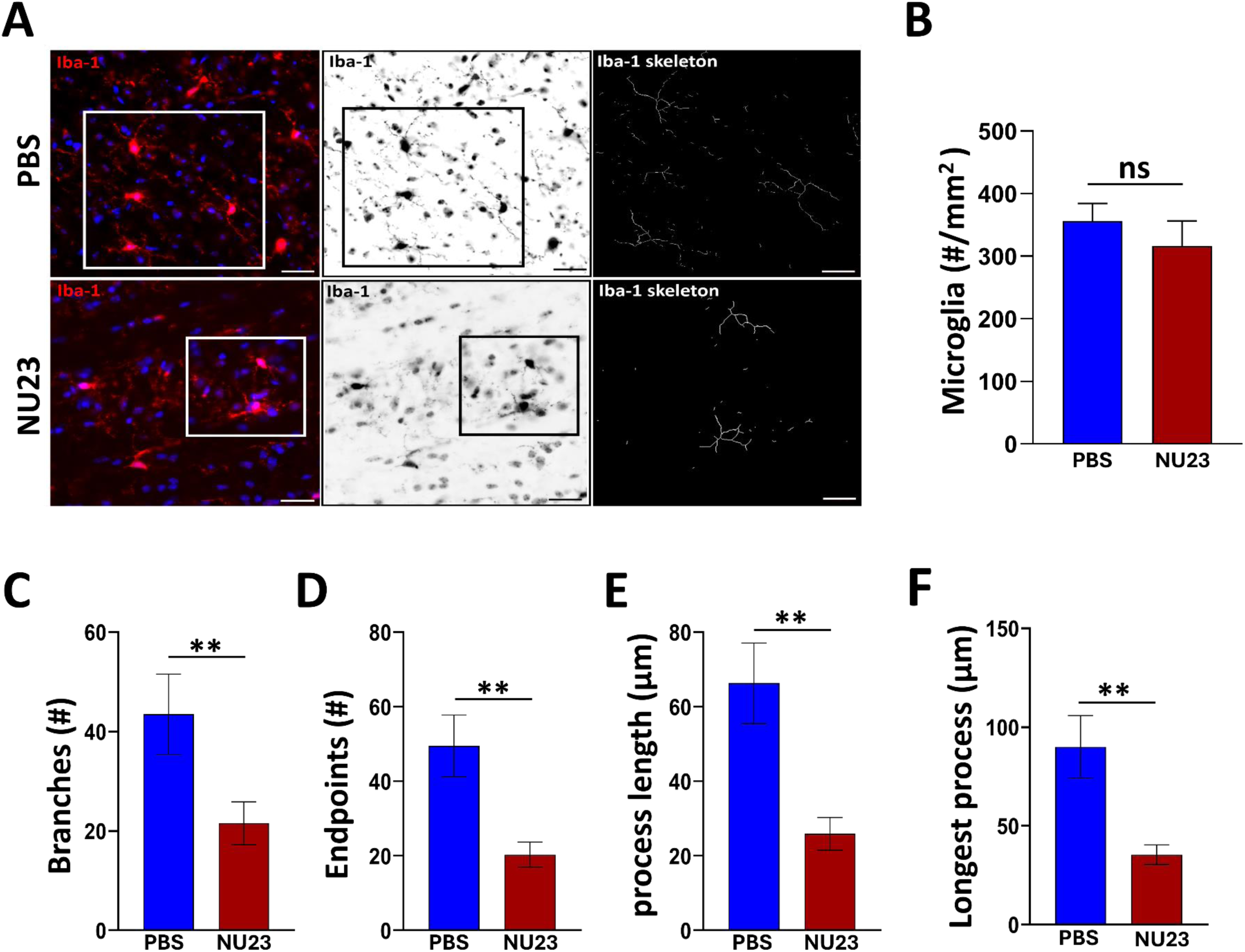
Post-UTI pelvic pain is associated with an activated microglial morphology in the prefrontal cortex. **(A)** Representative micrographs of Iba1-positive microglia in the prefrontal cortex of PBS– and NU23-instilled mice at 3 weeks post-instillation. Left, representative Iba1 immunofluorescence images (red) with DAPI nuclear counterstaining (blue). Middle, grayscale Iba1 images used for morphometric analysis. Right, skeletonized images generated for quantitative assessment of microglial branching complexity. Scale bar = 50 μm. **(B)**Quantification of the number of Iba1-positive microglia in the prefrontal cortex of PBS– and NU23-instilled mice. **(C–F)** Quantification of microglial morphology from skeletonized Iba1 images in PBS– and NU23-instilled mice. Data are presented as mean ± SEM. Statistical significance was determined using Welch’s t-test. *P < 0.05, **P < 0.01.

### Microglia from mice with chronic pelvic pain exhibit transcriptional signatures of activation

To define the molecular state of microglia associated with UPEC-induced chronic pelvic pain, we performed transcriptomic profiling of CD11b+ brain cells isolated three weeks after infection. Principal component analysis revealed clear separation between UPEC and control samples, indicating persistent transcriptional reprogramming following infection **(Fig. 3A)**. In many neuroinflammatory conditions, activated microglia downregulate homeostatic markers and transition toward a disease-associated phenotype **[38]**. In contrast, our RNA-seq data showed that microglia from NU23-instilled mice retained robust expression of canonical homeostatic microglial genes such as *Tmem119* and *P2ry12* **(Fig. S4)**. This suggests that the cells remain identifiable as resident microglia rather than undergoing a complete loss of microglial identity where homeostatic markers are lost while upregulating the expression of markers involved in activation such as inflammatory signaling. Differential gene expression analysis identified increased expression of chemokines (*Ccl2*, *Ccl6*, and *Ccl7*), NFΚB-associated genes (*Nffib2*), immediate early response genes (*Fosb* and *Jund*), and adhesion and motility genes (*Itga5, Lpxn*), together with genes involved in immune cell interactions and myeloid activation such as *Mpo*, *Ly6c1*, and *Ctse*. In contrast, pro-inflammatory cytokine transcripts such as *Tnf-α* and *Il-6* were not prominently induced **(Fig. 3B and 3C)**. Among the most significantly decreased genes were *Trpm3*, *Ig@p2*, *Vat1l*, *Clic6*, *Atp1b1*, *Cntn1*, *Tenm4*, and *Cx3cl1* which are broadly implicated in ion channel activity, growth factor signaling, vesicular trafficking, and cell adhesion processes that support microglial sensing of and communication with the neural environment **(Fig. 3B)**. Gene ontology analysis further demonstrated enrichment of pathways associated with leukocyte recruitment, immune regulation, lymphocyte chemotaxis, and TNF signaling **(Fig. 3D)**. Collectively, these findings indicate that post-UTI chronic pelvic pain is associated with persistent reactive microglial state characterized by enhanced reactivity and chemokine signaling.

**Figure 3.**
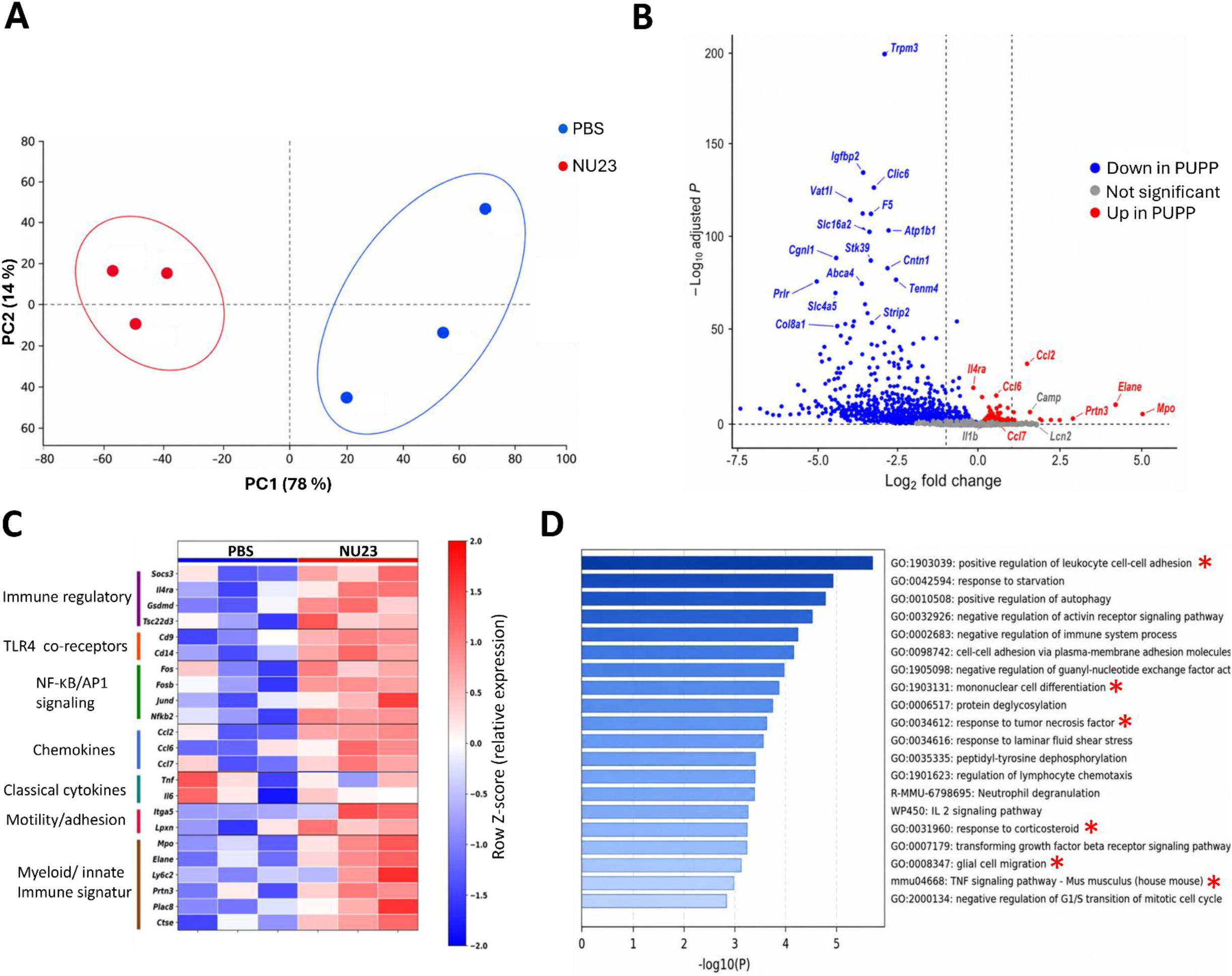
Transcriptomic profiling reveals microglial activation following infection with *E. coli* NU23. **(A)** Principal component analysis (PCA) of prefrontal cortex microglia isolated from control and NU23-instilled mice 3 weeks post-instillation, showing distinct transcriptional clustering between groups. **(B)** Volcano plot of differentially expressed genes in microglia from NU23-instilled mice relative to controls. Red and blue dots represent significantly upregulated and downregulated genes, respectively. Differential expression was defined as log₂ fold change > 1. **(C)** Heatmap of selected differentially expressed genes associated with inflammatory signaling, chemokine production, NFΚB activation, cell adhesion, migration, and brain-infiltrating leucocytes. Genes shown met an adjusted P < 0.05 threshold. Each column represents an individual biological replicate. **(D)** Gene Ontology and pathway enrichment analysis of genes upregulated in microglia from NU23-instilled mice, highlighting pathways related to immune regulation, inflammatory signaling, cell adhesion, and glial migration. RNA sequencing was performed on microglia isolated from the prefrontal cortex of control and NU23-instilled mice 3 weeks post-instillation (n = 3 mice per group).

### TLR4 drives microglial inflammatory signaling in vitro

To investigate the mechanisms by which TLR4 regulates microglial activation, BV-2 microglia were treated with a TLR4 antagonist prior to LPS stimulation. LPS induced robust cytokine production, whereas pharmacological inhibition of TLR4 significantly attenuated this response, indicating that TLR4 signaling is required for microglial inflammatory activation **(Fig. 4A)**. To directly assess the contribution of TLR4 to NFΚB signaling in microglia, we generated a stable BV-2 cell line expressing an NFΚB-responsive luciferase reporter. Consistent with the cytokine secretion data, LPS stimulation markedly increased NFΚB-dependent luciferase activity in BV-2 NFΚB reporter cells, whereas pharmacological inhibition of TLR4 significantly attenuated NFΚB activation **(Fig. 4B)**. Furthermore, LPS stimulation increased expression of the activation marker CD68, an effect that was largely prevented by TLR4 antagonism **(Fig. 4C and D)**. Together, these findings demonstrate that TLR4 promotes microglial activation through NFΚB and support a mechanistic role for TLR4 in driving the reactive microglial phenotype observed in vivo following infection with NU23.

**Figure 4.**
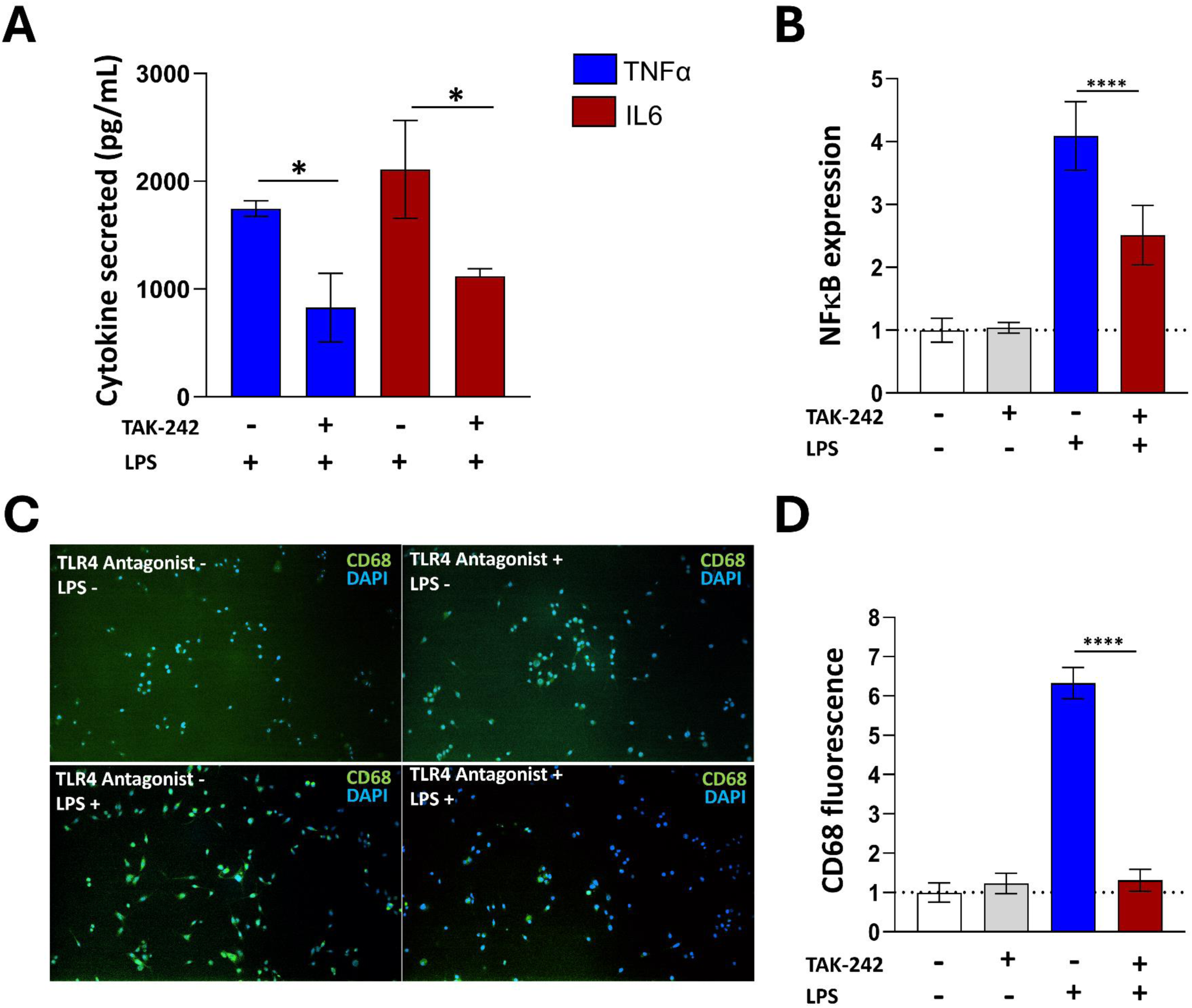
Pharmacological inhibition of TLR4 suppresses microglial inflammatory responses and NFΚB activation in vitro. **(A)** Secretion of TNF-α and IL-6 by BV-2 microglia following stimulation with LPS in the presence or absence of the TLR4 antagonist TAK-242. Cytokine concentrations were measured by ELISA. **(B)** NFΚB-dependent luciferase activity in BV-2 NFΚB reporter cells following stimulation with LPS in the presence or absence of TAK-242. **(C)** Representative immunofluorescence images of CD68 expression (green) in BV-2 microglia treated with vehicle or TAK-242 and stimulated with LPS as indicated. Nuclei were counterstained with DAPI (blue). **(D)** Quantification of CD68 fluorescence intensity in BV-2 microglia under the indicated treatment conditions. Data in panels A and B are presented as mean ± SEM from three independent experiments, each performed with three technical replicates per condition. CD68 fluorescence intensity in panel D was quantified from 20 fields obtained from three independent slides per experimental group. Statistical significance was determined using Welch’s t-test. P* < 0.05, P** < 0.01, P*** < 0.001, P**** < 0.0001.

### TLR4 signaling contributes to microglial activation and pelvic pain

To investigate the role of TLR4 in PUPP, we compared pelvic allodynia between wild-type (WT) and TLR4-deficient mice following NU23 infection. Deletion of TLR4 significantly attenuated the development of pelvic allodynia, indicating that TLR4 is required for the full expression of the pain phenotype **(Fig. 5A)**. To specifically assess the role of microglial TLR4, we generated microglia-specific *Tlr4* conditional knockout mice that carry the *Tmem119-Cre* transgene and homozygous for the floxed *Tlr4* allele. The Cre/loxP recombination was induced by tamoxifen prior to bladder instillation with NU23, and pain responses were evaluated 3 weeks post-instillation. Similar to systemic TLR4 deficiency, microglial TLR4 deletion markedly reduced pelvic allodynia, demonstrating that TLR4 signaling within microglia contributes substantially to post-infectious pelvic pain **(Fig. 5B)**. In WT mice, NU23 infection induced a reactive microglial phenotype characterized by reduced branching complexity, shortened processes, and a less ramified morphology. In contrast, microglia from TLR4 knockout mice retained a more homeostatic morphology **(Fig. 5C)**. Quantitative morphometric analyses showed increased process length, branch number, and overall cellular complexity in TLR4-deficient microglia relative to WT mice, indicating that loss of microglial TLR4 signaling prevents activation-associated structural morphological remodeling **(Fig. 5D-G)**. Importantly, preservation of a ramified microglial architecture in knockout mice coincided with a marked reduction in pelvic allodynia, linking microglial TLR4 signaling to both morphological activation and pain behavior. Together, these findings identify microglial TLR4 as a key regulator of microglial activation and post –UTI chronic pelvic pain.

**Figure 5.**
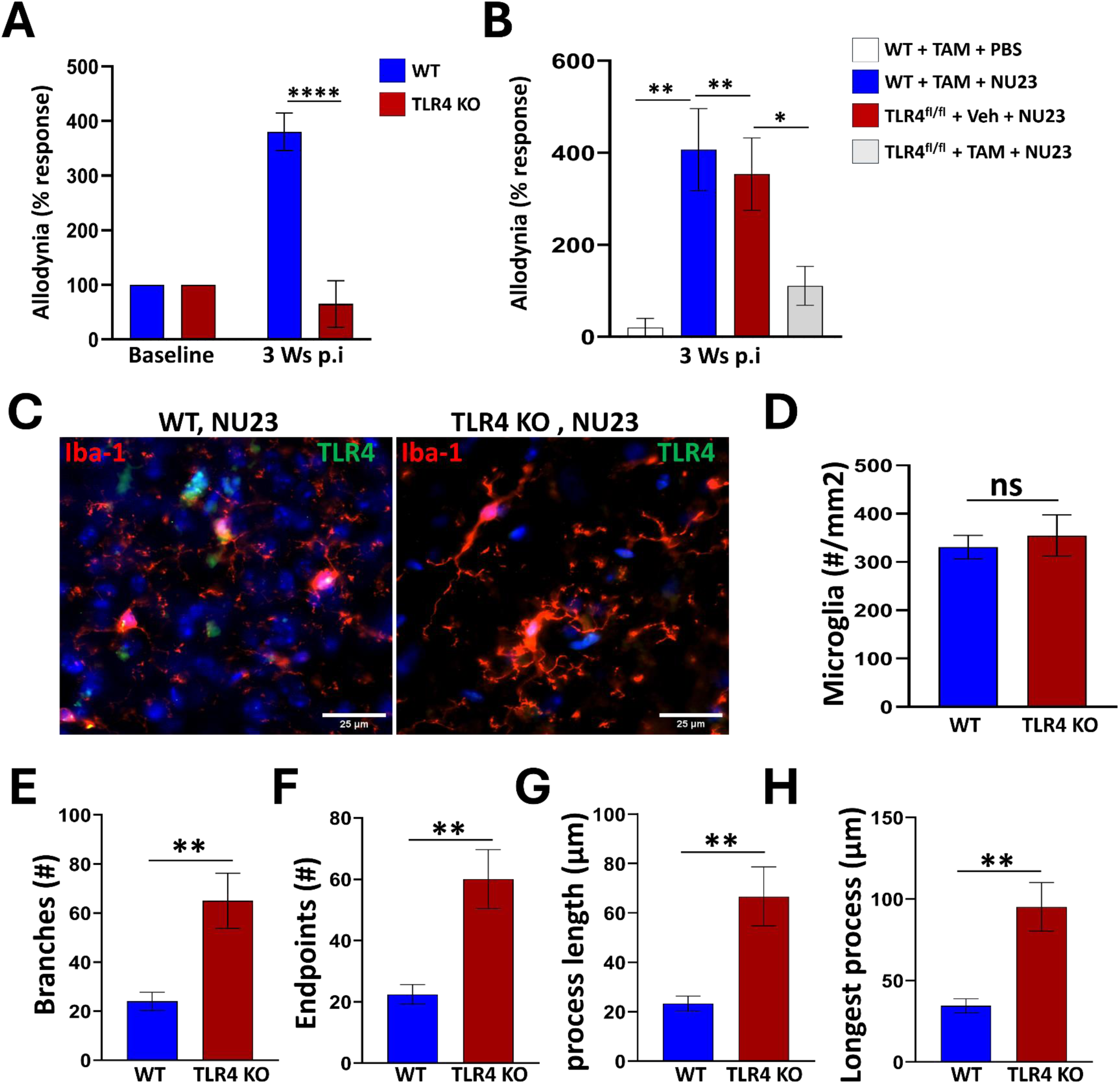
TLR4 contributes to post-UTI pelvic pain and microglia activation. **(A)** Pelvic allodynia assessed 3 weeks following PBS or NU23 instillation in wild-type and general TLR4 KO mice. **(B)** Pelvic allodynia assessed 3 weeks following PBS or NU23 instillation in control and microglia-specific TLR4 KO mice. **(C)** Representative immunofluorescence images of prefrontal cortex microglia labeled for Iba1 (red) and TLR4 (green) in WT and TLR4 KO mice. Nuclei were counterstained with DAPI (blue). Scale bar = 25 μm. **(D)** Quantification of Iba1-positive microglia in the prefrontal cortex of WT and TLR4 KO mice 3 weeks after instillation with NU23. **(E–H)** Quantification of microglial morphological parameters, including number of branches and endpoints, mean process length, and longest process length, in the prefrontal cortex of WT and TLR4 KO mice following NU23 instillation. Data are presented as mean ± SEM. Statistical significance was determined using Welch’s t-test. P* < 0.05, P** < 0.01, P*** < 0.001, P**** < 0.0001.

## Discussion

The mechanisms linking a prior history of UTI to the persistent symptoms of IC/BPS remain poorly understood. Here, we identify microglial TLR4 as a critical mediator of post-infectious chronic pelvic pain. Pharmacological depletion of microglia alleviated pelvic pain following UTI. In addition, both global and microglia-specific deletion of TLR4 attenuated microglial activation and prevented the development of chronic pelvic pain. Notably, microglial depletion did not improve urinary dysfunction or anxiety– and depression-like behaviors, indicating that microglia modulate the pain component of IC/BPS rather than all disease manifestations. Consistent with this observation, Griffith and colleagues, through the MAPP Research Network, demonstrated that pain and urinary symptom severity are dissociated in patients with urological chronic pelvic pain syndrome (UCPPS), supporting the concept that distinct biological mechanisms underlie different symptom domains of IC/BPS **[39]**.

Mechanistically, our transcriptomic data indicate that UTI induces a persistent microglia activation that sustain chronic pain by modulating neuron–glia communication and central sensitization. In contrast, persistent urinary dysfunction is likely maintained by bladder pathology and alterations in urothelial–afferent signaling, autonomic reflex pathways, and central micturition circuits **[40–42]**, processes that may be independent of microglia. Likewise, the lack of an effect of microglia ablation on anxiety– and depression-like behaviors suggests that the microglial changes induced by transient UTI alone are not sufficient to drive the widespread neuroimmune and synaptic alterations in corticolimbic circuits that are thought to underlie anxiety– and depression-like behaviors **[43,44]**.

Morphometric analyses of microglia demonstrated an activated amoeboid morphology in NU23-induced PUPP mice, consistent with chronic microglial activation. In addition, transcriptomic analyses of microglia isolated three weeks after infection showed that pro-inflammatory cytokines such *Tnf-α* and *Il-1β* were not significantly elevated. Instead, microglia exhibited increased expression of *Nffib2*, chemokines *Ccl2* and *Ccl5*, and immediate-early genes. These changes were accompanied by increased cell adhesion, cytoskeletal remodeling, and motility genes, suggesting a sustained shift toward a more reactive and dynamically interacting microglial phenotype rather than a cytokine-dominant inflammatory state. At three weeks post-UTI, cytokines secreted by microglia may have already peaked earlier and returned to baseline, while chemokine and stress-response programs persist, as observed in previous studies **[45, 46]**. The downregulated genes provide additional insight into the nature of the microglial response following NU23 infection. Many of the most strongly decreased genes were associated with membrane excitability and cell-cell communication. Such a transcriptional profile is consistent with a primed microglial state that may sustain chronic pain through persistent neuroimmune communication rather than overt cytokine-driven inflammation **[47,48]**.

Both global and microglia-specific TLR4 deficiency prevented the development of chronic pelvic pain following UTI, demonstrating that microglial TLR4 is a critical driver of post-infectious pain sensitization. These findings extend our previous work implicating peripheral TLR4 in chronic pelvic pain development **[28]** and complement clinical studies, including those from the MAPP Research Network, which have associated TLR4-mediated inflammatory responses with symptom severity in IC/BPS **[23, 26, 27, 49, 50]**. While these studies primarily implicate peripheral TLR4 signaling, our results identify microglial TLR4 as a central mechanism linking transient bladder infection to persistent pelvic pain.

Several clinical findings support a role for microglia in chronic pain. Positron emission tomography (PET) studies using translocator protein (TSPO) ligands have demonstrated increased glial activation in brain regions involved in pain processing in patients with chronic low back pain, fibromyalgia, and complex regional pain syndrome **[51, 52]**. In addition, elevated concentrations of inflammatory mediators have been reported in the cerebrospinal fluid of patients with chronic pain including fibromyalgia, neuropathic pain, and complex regional pain syndrome, suggesting persistent central immune activation **[53–55]**. Furthermore, clinical trials testing several drugs with microglia-modulating properties including minocycline, ibudilast, and low-dose naltrexone, have shown analgesic benefits in patients with chronic pain disorders other than IC/BPS **[56–60]**. Thus, our findings extend the forementioned studies by identifying microglia and their TLR4 as a potential therapeutic target in IC/BPS.

In conclusion, our findings demonstrate that microglial TLR4 is a key mediator of post-UTI chronic pelvic pain but plays a limited role in urinary dysfunction and anxiety/depression-like behaviors. Transient infection induced a persistent microglial activation characterized by increased chemokine and immediate-early gene expression despite minimal induction of canonical pro-inflammatory cytokines. These results support a model in which transient bladder infection drives long-lasting neuroimmune alterations within the CNS that sustain chronic pelvic pain after infection resolution and identify microglial TLR4 as a potential therapeutic target for IC/BPS.

### Ethical Approval Statement

All animals were maintained at the Center for Comparative Medicine at Northwestern University. Animal studies were performed under protocols approved by the Institutional Animal Care and Use Committee of Northwestern University.

### Author Contributions

H.J. and D.J.K. conceived and designed research; H.J and S.G performed experiments; H.J. analyzed data; H.J., A.J.S. and D.J.K. interpreted results of experiments; H.J. drafted manuscript; D.J.K. edited and revised manuscript; D.J.K. approved final version of manuscript.

### Author Approvals

All authors have seen and approved the manuscript. This manuscript hasn’t been accepted or published elsewhere.

### Funding

This work was supported by NIDDK 1R01DK134817.

