## Supplemental Material for "Microglial TLR4 Mediates Post-UTI Chronic Pelvic Pain"

* to whom correspondence should be addressed

312.908.1996

### Figure S1

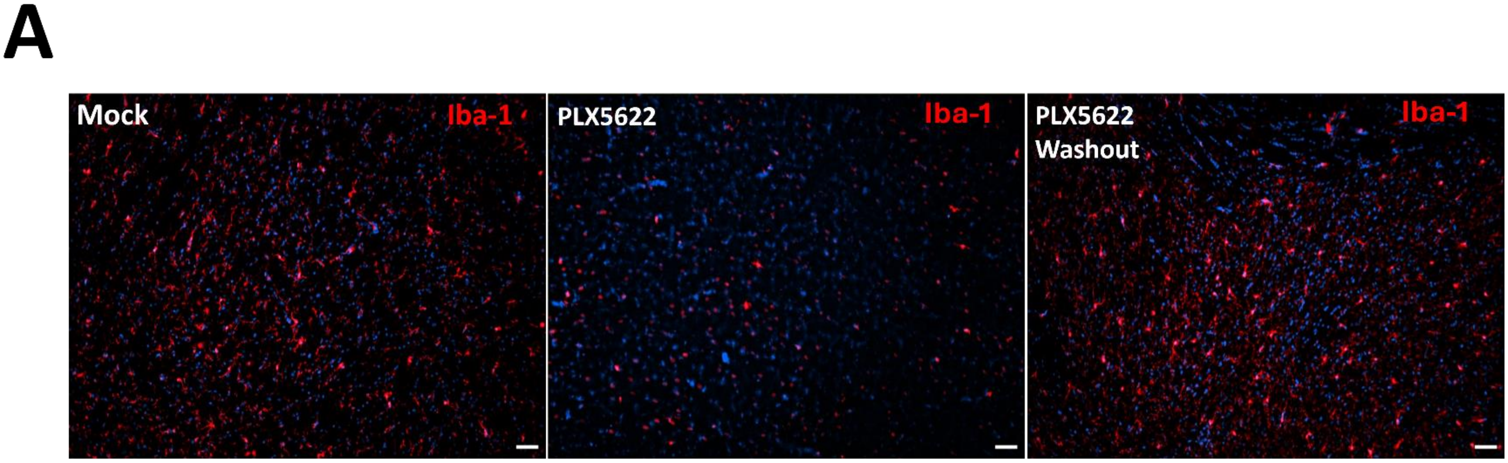


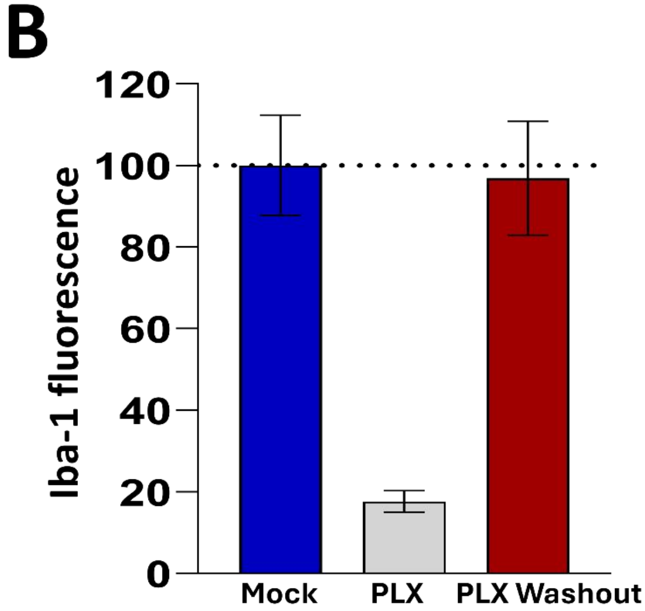


**Figure S1**. PLX5622 efficiently depletes microglia and allows repopulation following washout. **(A)** Representative Iba1 immunofluorescence images of the prefrontal cortex from mock-treated mice, PLX5622-treated mice, and mice following PLX5622 washout. Iba1-positive microglia are shown in red and nuclei are counterstained with DAPI (blue). Scale bars = 25 µm. **(B)** Quantification of the number of Iba1-positive microglia in the prefrontal cortex of mock-treated mice, PLX5622-treated mice, and mice following PLX5622 washout. Data are presented as mean ± SEM.

### Figure S2

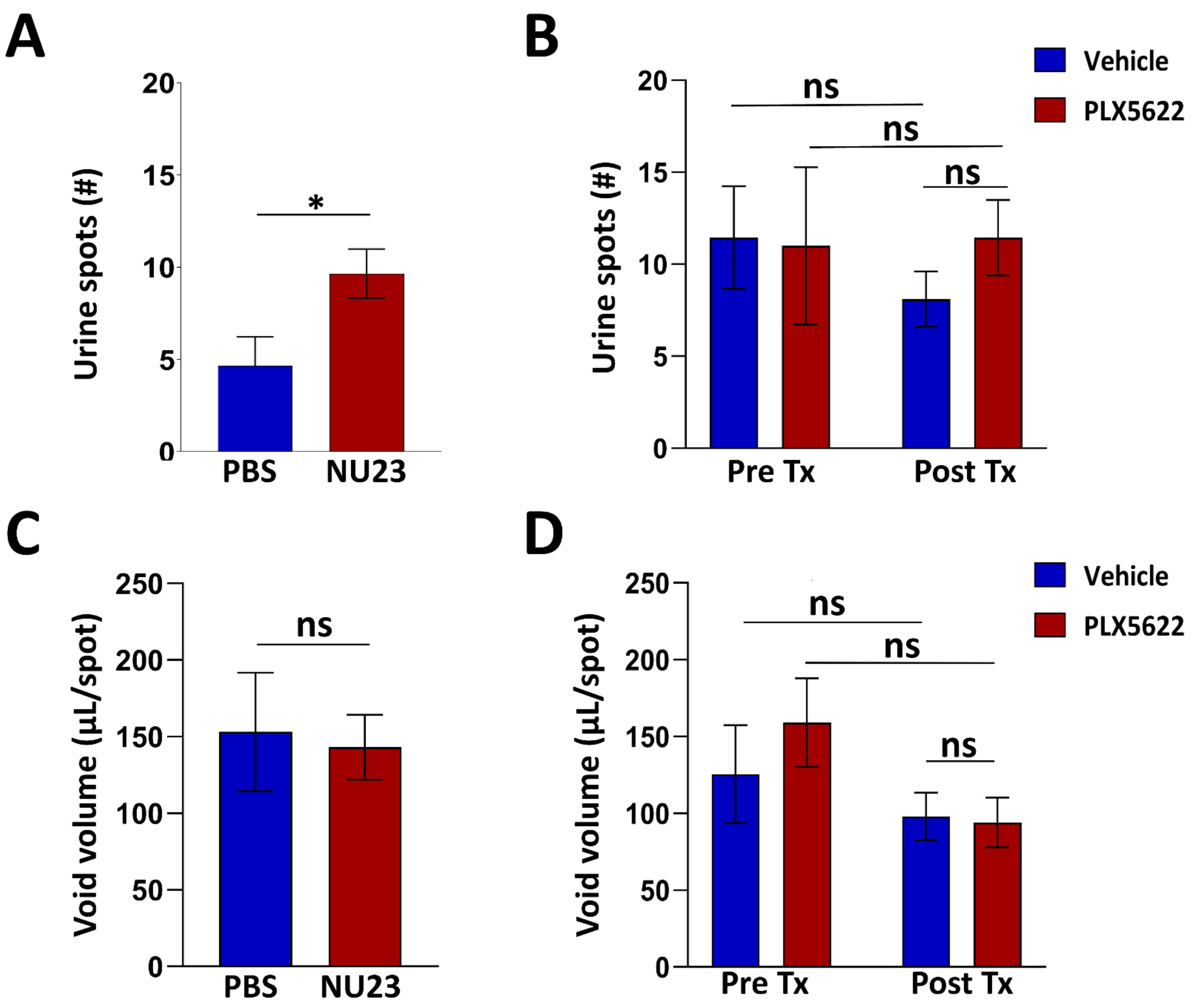


**Figure S2**. Microglial ablation does not alter voiding dysfunction in mice with post-UTI pelvic pain. **(A)** Number of urine spots measured by void spot assay in PBS- and NU23-instilled mice 3 weeks after instillation. **(B)** Number of urine spots measured by void spot assay in NU23-instilled mice before treatment (left) and following treatment with vehicle or PLX5622 (right). **(C)** Average urine volume per spot measured by void spot assay in PBS- and NU23-instilled mice 3 weeks after instillation. **(D)** Average urine volume per spot measured by void spot assay in NU23-instilled mice before treatment (left) and following treatment with vehicle or PLX5622 (right). Data are presented as mean ± SEM. Statistical significance was determined using Welch's t-test. $P^* < 0.05$.

### Figure S3

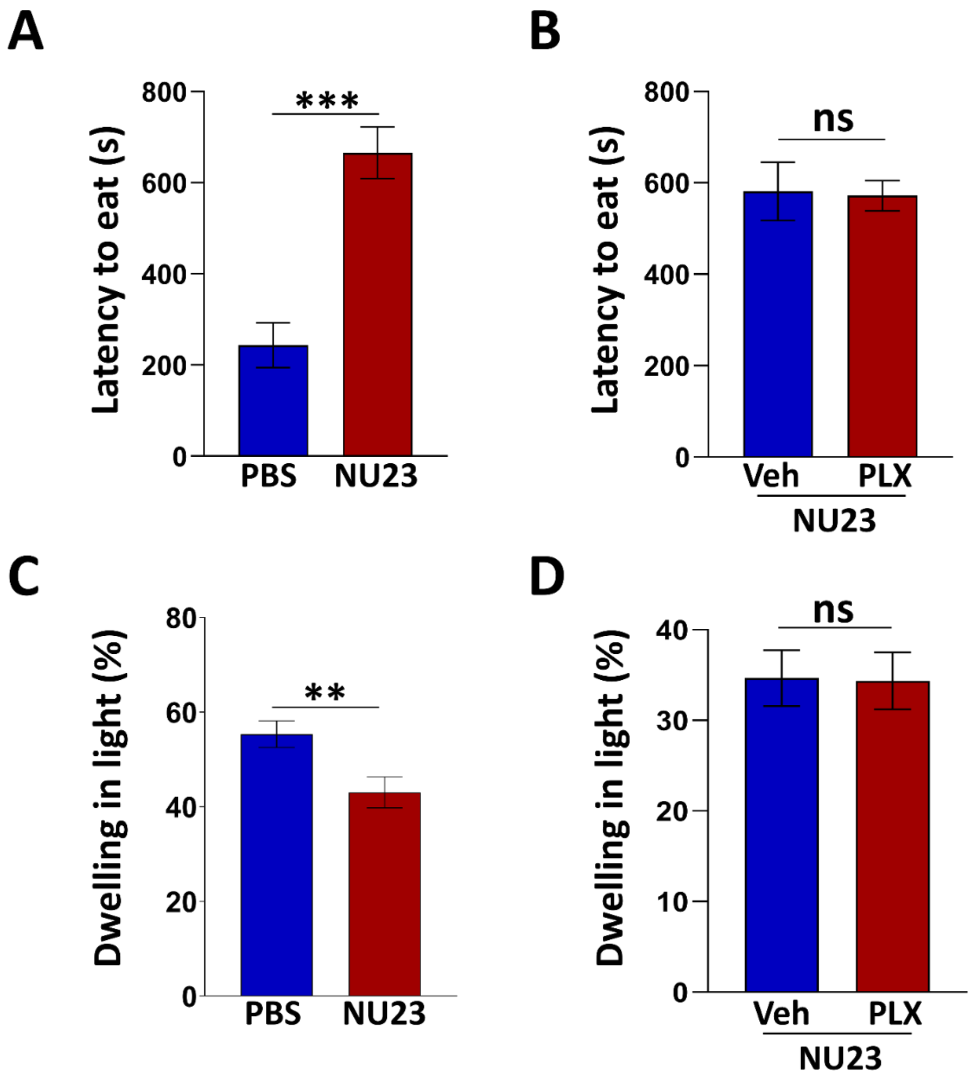


**Figure S3**. Microglial ablation does not improve anxiety- and depression-like behaviors in mice with post-UTI chronic pelvic pain. **(A)** Time spent in the light compartment during the dark-light box test in mice instilled with PBS or NU23. **(B)** Time spent in the light compartment during the dark-light box test in mice with PUPP treated with vehicle or PLX5622. **(C)** Latency to eat during the novelty-suppressed feeding test in NU23-instilled mice compared with PBS-instilled mice. **(D)** Latency to eat measured during the novelty-suppressed feeding test in NU23-instilled mice treated with vehicle or PLX5622. Data are presented as mean ± SEM. Statistical significance was determined using Welch's t-test. P* < 0.05, P** < 0.01, P*** < 0.001.

### Figure S4

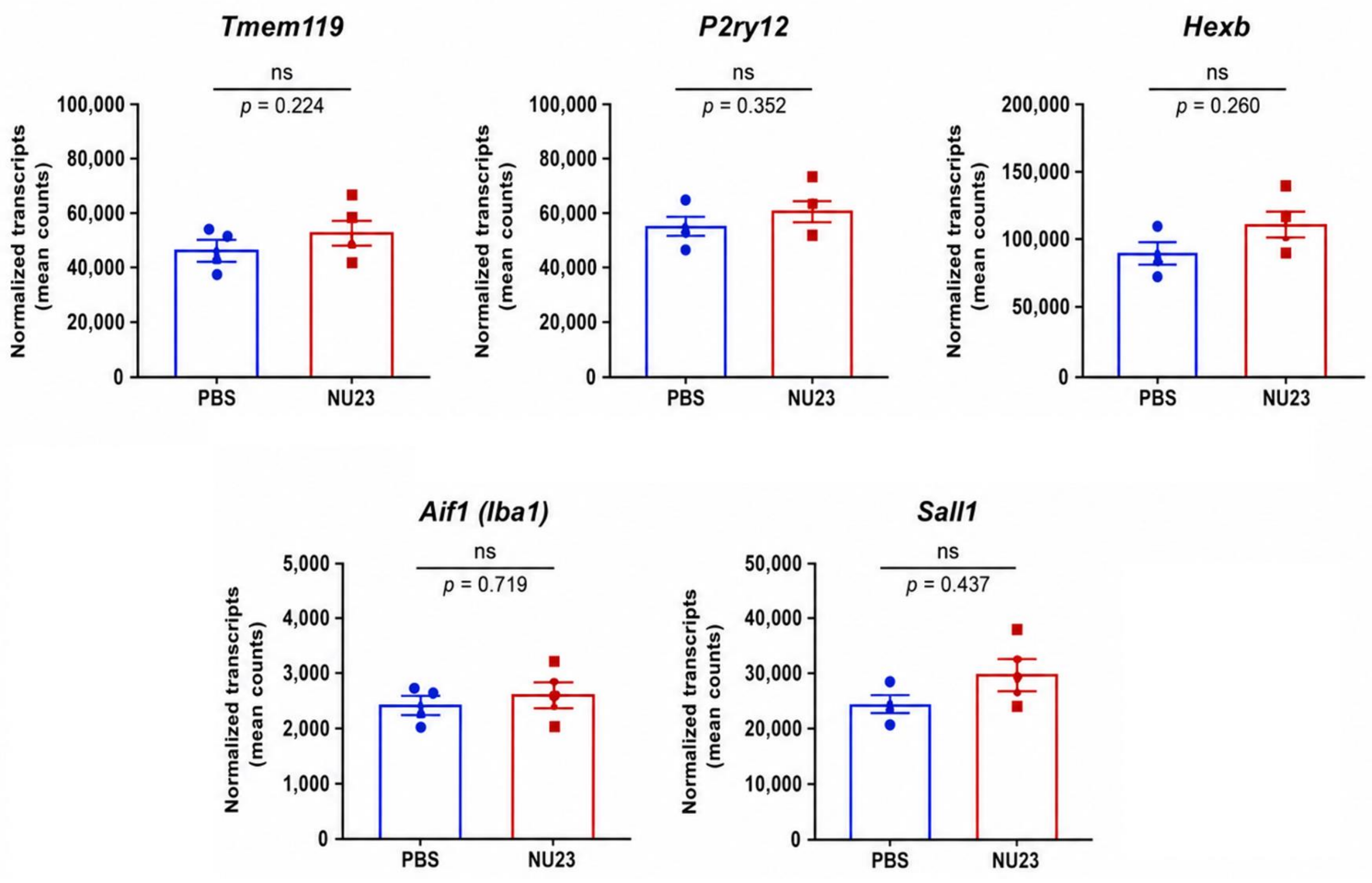


**Figure S4**. Homeostatic microglial gene expression following NU23 instillation. Normalized transcripts count for *Tmem119*, *P2ry12*, Hexb, *Aif1 (Iba1)*, and *Sall1* in PFC microglia isolated from mice 3 weeks after transurethral instillation of PBS or NU23. Individual biological replicates are shown together with mean ± SEM (n = 3 mice/group). Differential expression analysis was performed using DESeq2. No significant differences were detected between groups (ns, FDR ≥ 0.05).
